# Multi-Task Reinforcement Learning in Humans

**DOI:** 10.1101/815332

**Authors:** Momchil S. Tomov, Eric Schulz, Samuel J. Gershman

## Abstract

The ability to transfer knowledge across tasks and generalize to novel ones is an important hallmark of human intelligence. Yet not much is known about human multi-task reinforcement learning. We study participants’ behavior in a novel two-step decision making task with multiple features and changing reward functions. We compare their behavior to two state-of-the-art algorithms for multi-task reinforcement learning, one that maps previous policies and encountered features to new reward functions and one that approximates value functions across tasks, as well as to standard model-based and model-free algorithms. Across three exploratory experiments and a large preregistered experiment, our results provide strong evidence for a strategy that maps previously learned policies to novel scenarios. These results enrich our understanding of human reinforcement learning in complex environments with changing task demands.

---

Imagine standing at the threshold of your apartment on a Sunday morning (Figure 1A). If you are feeling hungry, you might choose to go to your local burger joint where they make terrific burgers. Conversely, if you are feeling groggy, you might instead go to your favorite coffee shop where they make the best espresso. Now imagine that, on a rare occasion, you are feeling both hungry and groggy. In that case, you might choose to go to a local diner where they make decent food and decent coffee. Even if you have only had food or coffee there up to that point, and would prefer the burger joint or the coffee shop over the diner for either of them, you will be able to extrapolate to the rare scenario in which you need both and realize that the diner is the best option. Finally, imagine that one day you decide to start reading fiction for fun, but would like to do it in a social setting. Given this new task, you might again choose to go to the coffee shop, even though you have never pursued this particular goal before.

**Figure 1.**
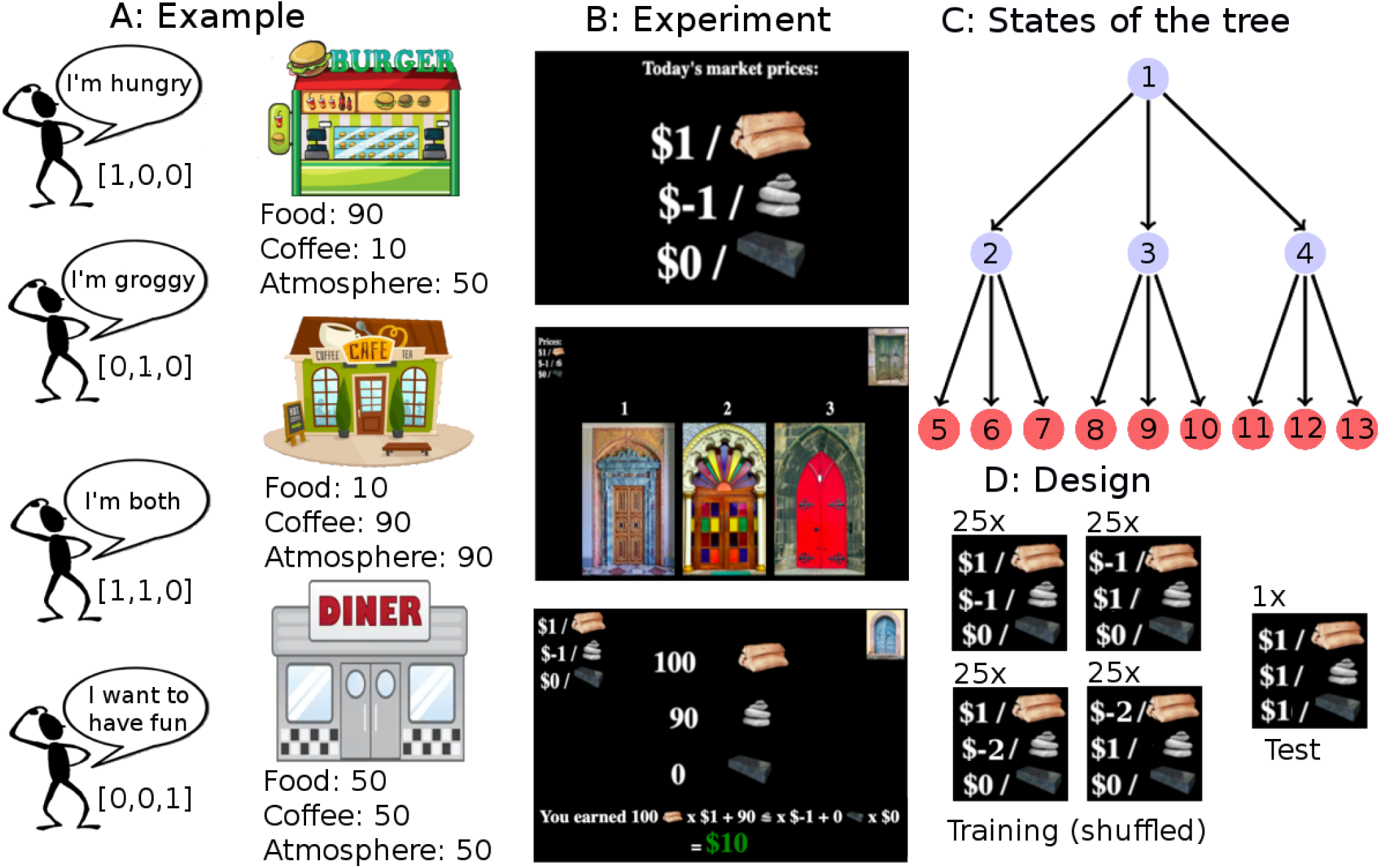
Overview of theoretical setup and experiments. **A:** Example of multi-task reinforcement learning. An agent who has found that going to the burger joint or coffee shop is a good way to gain rewards when hungry or groggy might have also encountered a diner before, which can be a viable option if the agent is both hungry and groggy. Moreover, the agent might have also learned that the coffee shop has good atmosphere and therefore could go there to have fun. **B:** Screenshots of the experiment. Participants were presented with a reward function on every trial, which corresponded to prices or costs they would receive/pay when encountering different resources (wood, stone, or iron) at the final state of the decision tree. Top: Participants saw the current price/costs for each resource at the beginning of each trial. This corresponds to the reward function, which can change across trials. Middle: Participants had to choose between three doors on every step of the decision making task, leading to 12 different doors in total. In the top left corner, they saw a reminder of the current reward function. In the top right corner, they saw which door they had chosen on the previous step. Bottom: After having gone through two different doors, participants saw the quantities of each resource at the new state, where the quantity directly corresponds to the features of the state and remained unchanged for each final state across all trials. **C:** Decision tree in which nodes correspond to different states. Although each state comes with a given set of features *φ*(⋅), we only set the features of the final states to values different from 0, corresponding to the encountered quantities of resources in our experiments. All transitions were deterministic in our experiments. **D:** Design of the reward function. During training, participants encountered reward functions with positive weights for one resource and negative weights for another resource. The third resource’s weight was always set to 0. At test, participants encountered a reward function with all weights set to 1.

In these examples, a single agent is performing multiple tasks, but the tasks all share some common structure. Humans are able to exploit this structure in the service of flexible behavior, without learning each task from scratch. The question of how humans achieve such flexibility has long preoccupied cognitive scientists.^1, 2^ In this paper, we study this question from the perspective of reinforcement learning (RL), where the computational goal is to choose actions that maximize long-term reward.^3^

One way to design a flexible RL algorithm is to use a model of the environment. If an agent knows the reward it expects to obtain in each state of the environment, along with the transition structure (how to get from one state to another), then it can use planning to identify the reward-maximizing route through the environment. This is the essence of *model-based* algorithms. Importantly for our purposes, these algorithms can adapt immediately when the reward or transition structure changes across tasks, without needing to learn from scratch. However, this flexibility comes at a cost: planning is computationally expensive. An agent acting in real-time does not have the luxury of re-planning every time a change in the environment is observed.

A different way to achieve flexibility is to directly learn a *value function* that maps states and actions to expected future rewards, without learning a model of the environment. This is the essence of *model-free* algorithms. If the learned function captures the shared structure across tasks, the agent can adapt to new tasks without learning from scratch. Since the learned function is usually cheap to compute, the agent can circumvent the costs of planning. The challenge is to design a task-dependent value function that can generalize effectively. If, for example, the agent learned a separate mapping for every task, then there would be no cross-task generalization, whereas at the other extreme, learning a common mapping across tasks would lead to catastrophic interference across tasks.

Recent work in computer science has grappled with this challenge, using new ideas about how to efficiently exploit the shared structure across tasks (see Methods for details). One idea, known as *universal value function approximators* (UVFAs)^4^, is to represent multiple value functions with a single function approximator that can generalize over both states and tasks. The key assumption underlying this idea is that values vary smoothly across tasks—a small change in task parameters yields a small change in values. If this assumption holds, then UVFAs can learn to generalize across tasks in the same way that function approximators learn to generalize across states within a task, simply by treating the task identifier as an input to the function approximator. In our running example, this is the kind of generalization that would lead you to choose the diner when you are both hungry and groggy; you already know that the diner is a reasonably good place to go when you are either hungry or groggy, and it is reasonable to expect that the value of the diner in this new task will be similar to the values learned for these previous tasks.

Another idea is to maintain a set of policies (for example, the optimal policies for tasks that you encounter frequently, such as feeling hungry or feeling groggy). These policies can be generalized to new tasks by choosing the action (across all cached policies) that leads to the states best satisfying the new task goals—an algorithm known as *generalized policy improvement*.^5, 6^ This generalization is made possible by learning a predictive representation of states (successor features, SF). The combination of successor features and generalized policy improvement is known as SF&GPI. In our running example, this is the kind of generalization that would lead you to the coffee shop when you want to read in a social setting; by imagining the features of the state where you end up by going to the coffee shop (tables, a relaxed atmosphere, intellectuals chatting with each other), you can figure out that the policy you usually follow when feeling groggy (the one that leads to the coffee shop) could serve you well in this new task.

Similarly to their single-task counterparts, UVFAs and SF&GPI exploit different kinds of structure to generalize to new tasks^7^: UVFAs rely on similarity in the value function across tasks, while SF&GPI relies on the shared structure of the environment. The two frameworks thus confer distinct and complementary advantages which would manifest as different generalization patterns in new tasks.

In our experiments, we test these predictions by asking participants to search for rewards in a two-step sequential decision making task. At the final states of the decision tree, participants encounter different quantities of resources (the features for each state). At the start of each trial, participants see the values of the different features for that trial. This induces a reward function over the final state of the tree, with states with more valuable features delivering greater overall rewards. Thus, the same state can deliver a large reward on one trial but a small or even negative reward on another trial. After being trained on multiple such reward functions (or *tasks*), participants encounter a test trial with a new reward function we designed to discriminate between the two multi-task reinforcement learning frameworks as well as standard single-task reinforcement learning approaches. Across three experiments, we find strong evidence for the SF&GPI strategy. We then preregistered our previous design and ran a large-scale replication study, again finding evidence for the SF&GPI strategy. We conclude that participants in our studies solve multi-task reinforcement learning problems by evaluating previously learned policies given the current reward function and the features of the different states.

## Results

Participants played a two-step decision making task (Fig. 1B). On each trial of this task, participants saw the prices and costs for three different resources: stone, wood, and iron. Participants then could pick between three different doors which would lead them to another room, where they again saw a set of three different doors, leading to 9 final states in total (Fig. 1C). Each of the final states was a room that contained different quantities of the resources, which were then multiplied by their prices/costs and added together, leading to participants’ reward on that specific trial. The transitions between rooms were deterministic (i.e., choosing a particular door always led participants to the same next room). There were 13 rooms in total, corresponding to the different states of the decision tree (Fig. 1C).

The prices and costs of the different resources on a given trial can be interpreted as the weight vector **w** of the multi-task RL problem; changing the weights corresponds to changing the task. We use this property of our task to train participants on one weight vector and then test them on a different weight vector. The weights during the training were the same throughout all experiments. Specifically, the set of training weights for the first 100 trials was **w**_train_ = {[1, −1, 0], [−1, 1, 0], [1, −2, 0], [−2, 1, 0]}, where each participant experienced each set of weights 25 times in random order and each resource was randomly assigned to one of the three weights for each participant. The weights on the 101st trial, which was the crucial test for our models’ predictions, were set to **w**_test_ = [1, 1, 1] (i.e., a novel task in which all resources were equally rewarding).

In all experiments, the interesting final states are 6, 7, 9, and 12. Much like the burger joint from our running example, state 6 always has a substantial amount of the first feature, making it the optimal state for tasks [1, −1, 0] and [1, −2, 0]. Much like the coffee shop from our running example, state 12 always has a substantial amount of the second feature, making it the optimal state for tasks [−1, 1, 0] and [−2, 1, 0]. Like the diner, state 9 has both the first and the second feature, however the negative task weights mean that it is never rewarding during training. Yet state 9 does deliver a substantial reward on the test task [1, 1, 1], which UVFAs can extrapolate. Importantly, single-task approaches such as model-free value function approximation would never choose state 9, as it is not rewarding during training. SF&GPI would also not choose state 9 for the same reason, since it is unlikely that a policy will be learned leading to state 9. Also like the coffee shop, state 12 additionally has substantial amount of the third resource, meaning that it delivers a high reward on the test task [1, 1, 1] as well. Assuming that training induces policies leading to states 6 and 12, SF&GPI will identify state 12 as the best choice on the test task. UVFAs are unlikely to choose state 12 since they learn to ignore the third feature as it is irrelevant during training, a crucial aspect of function approximation. Additionally, because the weights of the training features alternate between positive and negative, a model-free learner would simply converge on a state that consistently delivers a reward of 0. Finally, we always designate state 7 as the single optimal state on the test task, predicted by a model-based learner that knows the full structure of the environment, but not by the other models.

### Experiment 1

In the first experiment, participants (*N* = 226, mean age=35.2, SD=10.3, 92 females) were trained on four different weights (see Fig 2A), in which the first and second weight always showed diverging directions and the third weight was set to 0 on every training trial. For example, if on one trial iron had a value of $1, then wood could have had a negative value of $-1, and stone a value of $0. Participants were then trained on 100 trials of these weights, experiencing 25 trials for each of the different weights, assigned at random. This training regime favored the development of two optimal policies: one leading to state 6 for tasks [1, −1, 0] and [1, −2, 0], and one leading to state 12 for tasks [−1, 1, 0] and [−2, 1, 0]. On the final 101st trial, the prices of the different resources changed and all of them had an equal value of 1.

**Figure 2.**
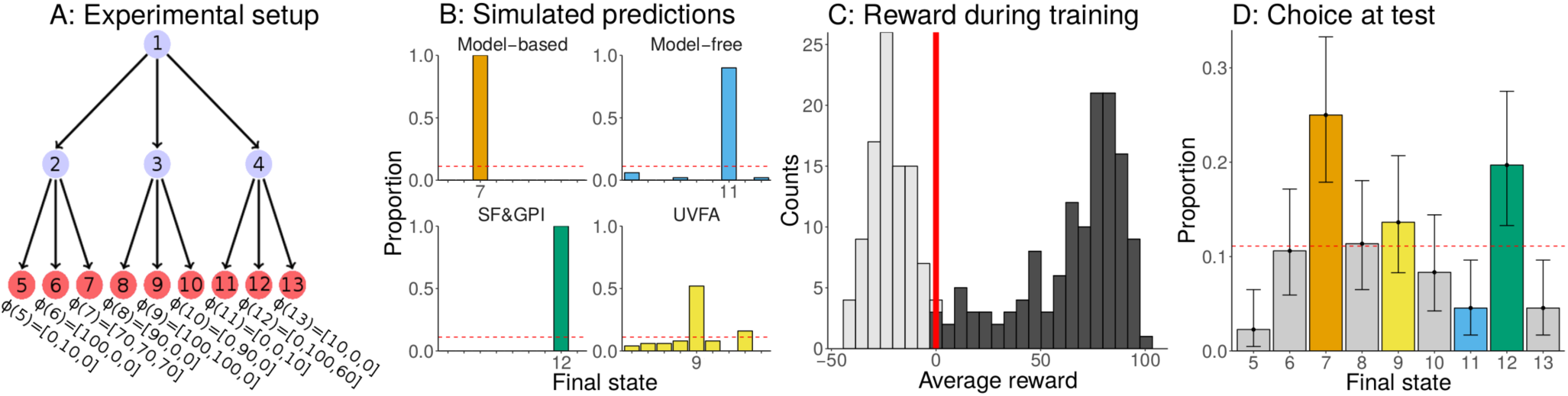
Overview and results of Experiment 1. **A:** Experimental setup. Participants are trained on the set of weights **w**_train_ = {[1, −1, 0], [−1, 1, 0], [1, −2, 0], [−2, 1, 0]} and tested on the weights **w**_test_ = [1, 1, 1]. The features for each final state are shown below the tree. **B:** Predictions of the different models. Predictions were derived by simulating models in our task given the training weights and then registering their decisions given the weights of the test trial. This simulation was repeated 100 times for each model and the proportions of choosing the different target states was tracked. **C:** Distribution of average reward obtained by participants during the training trials. Participants were split into a group that accumulated less than 0 points (gray) and a group that accumulated more than 0 points (black), which we analyzed further. Red vertical line marks the threshold of 0. **D:** Participants’ choices given the new weights **w**_test_ on the test trial. Choices are colored by the simulated model predictions. Error bars show the 95% confidence interval of the mean based on an exact binomial test. Dashed line indicates chance responding.

Importantly, we set the features of the final states to values that discriminated as much as possible between UVFAs, SF&GPI, and standard model-based and model-free reinforcement learning algorithms. In particular, the model-based algorithms predict that participants should go to state 7 with features *φ*(7) = [70, 70, 70], since given the new weights of **w** = [1, 1, 1] this state would produce the maximum possible reward of 210. Model-free reinforcement learning predicts that participants would go to state 11 with features *φ*(11) = [0, 0, 10], because that state led to an average reward of 0 and was thus the single best state during training, under the assumptions that the weights are ignored completely. UVFAs predict that participants would go to state 9 with features *φ*(9) = [100, 100, 0]. This is because this state maximizes the reward based on the first two weights, which UVFAs can extrapolate due to the smoothness of the training tasks, even though this state was never rewarding during training. Furthermore, UVFAs learn to ignore the third feature because its weight had been set to 0 throughout training. Finally, SF&GPI predicts that participants would go to state 12 with features *φ*(12) = [0, 100, 60], because it is the highest value state that the optimal training policies lead to. We verified these predictions by simulating the behavior of each model for 100 times in our task and calculating the probability with which each model chose the different final states on the 101st trial (Fig. 2B).

Next, we looked at participants’ performance in our task. Because the distribution of average rewards during the training trials was bimodal (see Fig. 2C, see Supplementary Information for further analyses), we only analyzed participants who gained an average reward of higher than 0 during the training rounds. This led to the exclusion of 94 participants. We then analyzed the remaining 132 participants’ decisions during the test trial (Fig. 2D). Thirty-three participants chose the final state 7 during the test trial. This state was predicted by a purely model-based learner, since it was the state leading to the highest overall reward. The proportion of participants choosing this state was significantly above the chance level of *p* = 1/9 (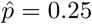, one-sided exact binomial test: *p* = 5.98 × 10^*−*6^, *BF* = 2359). The second most frequently chosen state was the final state 12, which was predicted by SF&GPI and chosen by 26 participants. This proportion was also significantly higher than chance (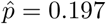, binomial test: *p* = .003, *BF* = 11.2). There was no significant difference between the two most frequently chosen options (*χ*^2^(1, 59) = 0.61, *p* = 0.435, *BF* = 0.45). None of the other states were chosen significantly more frequently than chance (all *p* > .05, max-*BF* = 0.43).

### Experiment 2

In Experiment 1, most participants chose the path that was predicted by a model-based planner, while the path predicted by SF&GPI was the second most frequently chosen path. However, the model-based path contained features that all had the same value. This means that in the test task, where each resource had the same value, the best state contained each of the resources in equal proportions. This might have helped people to choose the model-based path based on perceptually matching equal weights to a state with equal feature values. In Experiment 2, we attempted to overcome the effect of a matching between equal weights and features, by changing the features of the model-based path. Furthermore, we made the task more difficult by adding nuisance features that were not essential for the models’ optimal policies, but which made purely model-based planning computationally more demanding.

Participants (*N* = 202, mean age=37.6, SD=13.3, 85 females) participated in a similar task as in Experiment 1. However, we changed the feature values to create even stronger predictions about participants’ behavior (Fig. 3A). We increased the overall mean of all individual final states to create larger differences between them, given a particular weight. We also used more unique feature values in general in order to make the task harder and to not allow for chunking of feature values across states. We also changed the feature values of the target nodes to rule out competing explanations of the observed effects. First, we changed the features of state 7 to *φ*(7) = [70, 70, 170]. This was still the best possible state given the weights at test of **w** = [1, 1, 1], but did not allow for simple perceptual matching of equal weights to equal features. Furthermore, we changed the features of node 12 to *φ*(12) = [0, 100, 160] to make the attraction of the previously unrewarded weight even higher and create a stronger difference between the SF&GPI and all other models. Finally, we increased the two features of the previously rewarded weights for the state that was predicted by UVFAs to *φ*(9) = [150, 150, 0]. We verified these predictions by simulating 100 runs of all models in our task and calculating the proportion of final states they chose during the test trial (Fig 3B).

**Figure 3.**
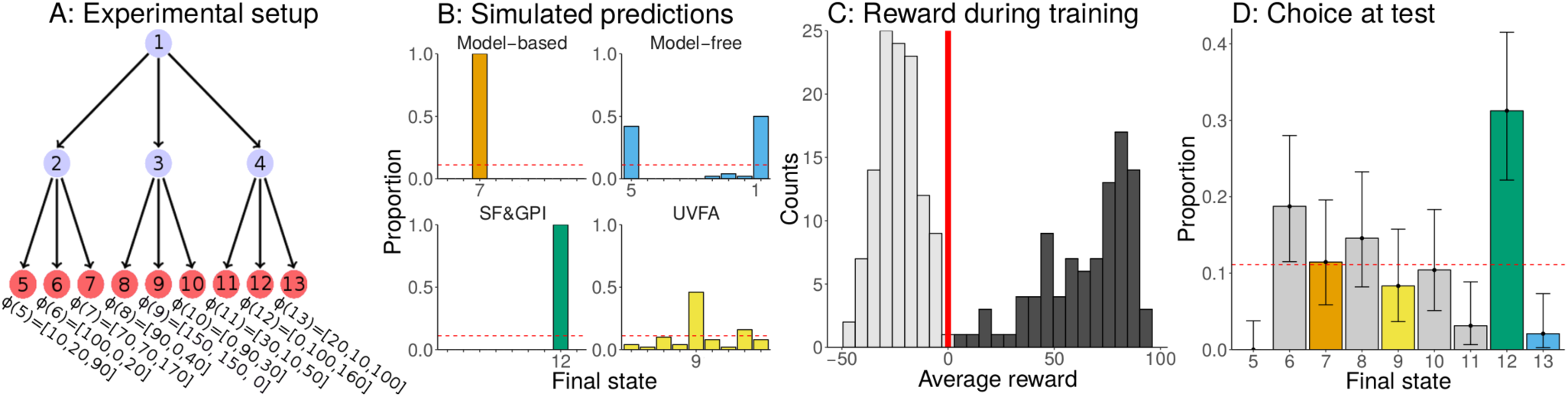
Overview and results of Experiment 2. **A:** Experimental setup. Participants are trained on the set of weights **w**_train_ = {[1, −1, 0], [−1, 1, 0], [1, −2, 0], [−2, 1, 0]} and tested on the weights **w**_test_ = [1, 1, 1]. The features for each final state are shown below the tree. **B:** Predictions of the different models. Predictions were derived by simulating models in our task given the training weights and then registering their decisions on the weight of the test trial. This simulation was repeated 100 times for each model and the proportions of choosing the different target states was tracked. **C:** Distribution of average reward obtained by participants during the training trials. Participants were split into a group that accumulated less than 0 points (gray) and a group that accumulated more than 0 points (black), which we analyzed further. Red vertical line marks the threshold of 0. **D:** Participants’ choices given the new weights in the test trial. Choices are colored by the simulated model predictions. Error bars show the 95% confidence interval of the mean based on an exact binomial test. Dashed line indicates chance responding.

As in Experiment 1, the performance distribution during training was bimodal (Fig. 2C). Thus, we again only looked at the 96 participants who earned an average reward higher than 0 and excluded the other 106 participants. The most frequently chosen path of these 96 participants was the one that led to state 12 (Fig. 2D), which was predicted by SF&GPI. In total, 30 of the 96 participants chose this state. This was higher than what would be expected at the chance level of *p* = 1/9 (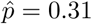, binomial test: *p* = 2.98 × 10^*−*6^, *BF* = 99837). Only 7 out of 96 participants chose the path that was predicted by a fully model-based planner. This was not significantly different from chance (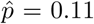, binomial test: *p* = .65, *BF* = 0.4) and also significantly lower than the number of people choosing the path predicted by SF&GPI (*χ*^2^(1, 41) = 7.90, *p* = .004, *BF* = 18.7). None of the remaining states that were predicted by the other models was chosen more frequently than what would be expected by chance (all *p* > .05, max-*BF* = 0.3).

Summarizing the results of Experiment 2: changing the features of the state predicted by a model-based reinforcement learning algorithm, and making the task more difficult in general, caused more participants to choose the state that was predicted by SF&GPI.

### Experiment 3

Since we changed multiple aspects of Experiment 1 in Experiment 2, we examined the robustness of our results by only changing fewer characteristics of the design applied in Experiment 1. In Experiment 3, we used a similar range of feature values as in Experiment 1, avoided the additional use of nuisance features, and chose the features of the final states so as to make as clear predictions as possible (Fig 4A).

**Figure 4.**
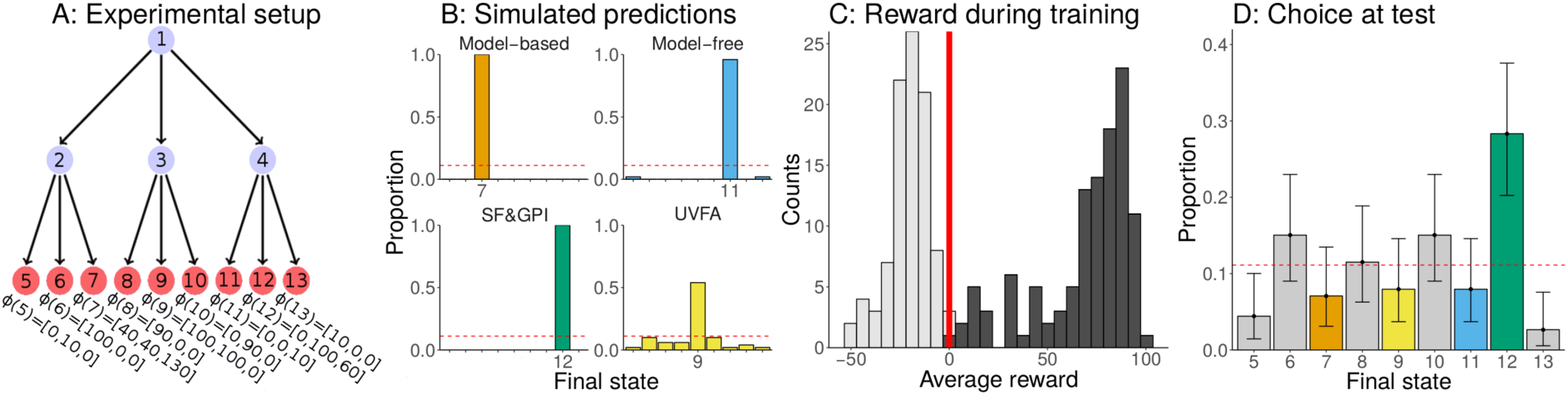
Overview and results of Experiment 3. **A:** Experimental setup. Participants are trained on the weights **w**_train_ = {[1, −1, 0], [−1, 1, 0], [1, −2, 0], [−2, 1, 0]} and tested on the weight **w**_test_ = [1, 1, 1]. The features for each final state are shown below the tree. **B:** Predictions of the different models. Predictions were derived by simulating models in our task given the training weights and then registering their decisions on the weight of the test trial. This simulation was repeated 100 times for each model and the proportions of choosing the different target states was tracked. **C:** Distribution of average reward obtained by participants during the training trials. Participants were split into a group that accumulated less than 0 points (gray) and a group that accumulated more than 0 points (black), which we analyzed further. Red vertical line marks the threshold of 0. **D:** Participants’ choices given the new weights in the test trial. Choices are colored by the simulated model predictions. Error bars show the 95% confidence interval of the mean based on an exact binomial test. Dashed line indicates chance responding.

**Figure 5.**
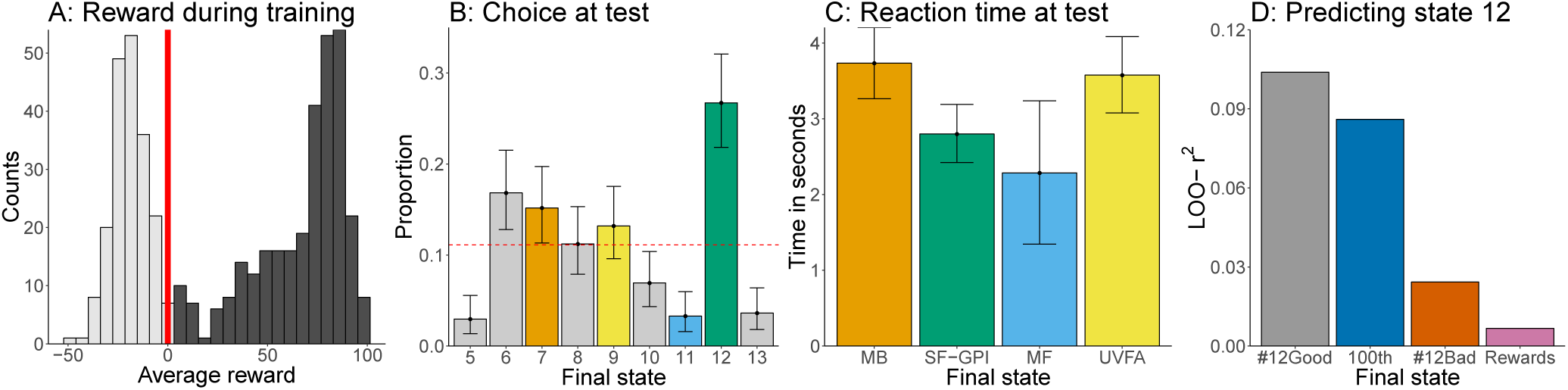
Results of preregistered Experiment 4. **A:** Distribution of average reward obtained by participants during the training trials. Participants were split into a group that accumulated less than 0 points (gray) and a group that accumulated more than 0 points (black), which we analyzed further. Red vertical line marks the threshold of 0. **B:** Participants’ choices given the new weights in the test trial. Choices are colored by the simulated model predictions. Error bars show the 95% confidence interval of the mean based on an exact binomial test. Dashed line indicates chance responding. **C:** Mean reaction times on the 101st trial in dependency of participants’ choices in seconds. Reaction times larger than 10 seconds have been removed. Error bars represent the 95% confidence interval of the mean. **D:** Bayesian model fit of different variables predicting participants choosing state 12 on the 101st trial. All values show the standardized (based on chance performance) approximate leave-one-out cross-validation error (LOO) based on the posterior likelihood of a logistic model regressing a variable onto a dependent variable indicating whether or not participants chose state 12. The regressors were how often participants correctly chose state 12 during training (#12Good), whether or not they chose state 12 on the 100th trial (100th), how often they incorrectly chose state 12 during training (#12Bad), and their average reward during training (Rewards).

This time, a model-based agent would again choose state 7 with *φ*(7) = [40, 40, 130], leading to the highest rewards given the test weights of **w** = [1, 1, 1]. A purely model-free agent would choose state 11 with *φ*(11) = [0, 0, 11], since that state on average leads to a reward of 0 during the training trials. UVFAs predict that people should choose state 9 with *φ*(9) = [100, 100, 0] by extrapolating from their experience with the training tasks. Finally, SF&GPI would pick state 12 with *φ*(12) = [0, 100, 60] because it is the best possible state to choose from the previously rewarding states, given the novel task. We validated these predictions by simulating 100 runs for each model in our task (Fig 4B).

As in the previous experiments, participants (*N* = 200, mean age=35.8, SD=11.2, 89 female) exhibited bimodal performance distributions during the training task was bimodal, and we therefore removed participants with an average reward below 0. This led to the exclusion of 87 of the 200 participants. Thirty-two of the remaining 113 participants chose the path that was predicted by SF&GPI. This state was the most frequently chosen state, significantly more than what would be expected under the chance level of *p* = 1/9 (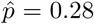, binomial test: *p* = 4.4 × 10^*−*7^, *BF* = 24902). The final state predicted by a model-based learner was only chosen by 8 out of 113 participants, which was not greater than chance (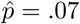, exact binomial test: *p* = .94, *BF* = 0.96) and significantly smaller than the number of people choosing the state predicted by SF&GPI (*χ*^2^(40, 1) = 13.23. *p* < .001, *BF* = 274). None of the other states was chosen significantly more often than chance (all *p* > .05, max-*BF* = 0.63).

Summarizing the results of Experiment 3: in a simpler setup of our two-step multi-task reinforcement learning task, we found further evidence for SF&GPI, since more people ended up choosing the state predicted by this model than by any alternative model.

### Experiment 4

All of the previous experiments produced at least partial evidence in favor of SF&GPI. However, given the bimodality of participants’ performance, we ended up excluding around 40% of participants in each experiment. Thus, to confirm the reproducibility of our results, we conducted a large preregistered replication of Experiment 3. Our preregistration included the exclusion criterion of removing participants with an average reward during training trials less than 0. We also committed ourselves to a sampling plan of collecting 500 participants in total (i.e., more than twice as many as in our previous experiments). Finally, we preregistered the following three hypotheses: (1) state 12 will be the most frequently chosen state overall, as predicted by SF&GPI; (2) state 12 will be chosen more frequently than chance (predicted *BF* of higher than 10); and (3) state 12 will be more frequently chosen than state 7 (predicted *BF* of higher than 10), because more people will follow the predictions of SF&GPI than those of a model-based planner.

Following our sampling plan, we recruited 500 subjects in total (mean age=34.66, SD=9.11, 214 females). As preregistered, we excluded all participants who did not achieve an average reward higher than 0 during the training trials. This led to the exclusion of 197 participants in total. Eighty-one of the remaining 303 participants chose the final state 12 on the 101st trial. Thus, state 12 was the the most frequently chosen state during the test trial, confirming our first preregistered hypothesis. The proportion of participants choosing states 12 was significantly higher than what would have been expected under the chance level of *p* = 1/9 (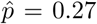, binomial test: *p* = 4.38 × 10^*−*14^, *BF* = 1.11 × 10^11^), confirming our second preregistered hypothesis. Finally, we assessed how many participants ended up choosing state 7, i.e. the state predicted by a model-based planner. Forty-six participants choose the final state 7, which was significantly more than chance (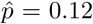, binomial test: *p* = .02, *BF* = 1.8). However, significantly more people chose the state corresponding to the SF&GPI prediction than the state corresponding to the model-based algorithm’s prediction (*χ*^2^(127, 1) = 9.10, *BF* = 22.4). This result confirmed our third and final preregistered hypothesis.

Because the final data set of Experiment 4 was much larger than our previous data sets, this allowed us to further investigate signatures of the SF&GPI model by exploring three additional analyses.

First, we looked at the proportion of participants who ended up choosing state 6 on the 101st trial. Although none of our models predicted this state *a priori*, it was frequently the best state (on half of all trials) during the training trials. Put differently, choosing state 6 was another policy participants could have learned during training. Since SF&GPI predicts that participants choose the best of the previously learned policies, this means that they should also prefer state 12 over state 6. In total, fifty-one participants ended up choosing state 6, a significantly larger proportion than chance (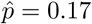, binomial test: *p* = .001, *BF* = 12.7). However, the proportion of participants choosing this state was significantly smaller than the number of participants choosing state 12 (*χ*(1, 132)^2^ = 6.37, *p* = .006, *BF* = 5.6), ruling out the possibility that participants chose one of the previously rewarded policies at random.

For the second exploratory analysis, we looked at participants’ reaction times on the 101st trial. Since SF&GPI simply re-evaluates previously encountered policies and features given the new reward function, this leads to the prediction that participants who choose in correspondence with the SF&GPI prediction (state 12) should do so faster than participants who choose in line with the model-based predictions (state 7). We therefore calculated participants’ mean reaction times on the 101st trial in dependency of their choices on that trial. We removed extremely large reaction times (above 10 seconds) before conducting this analysis. Comparing the different groups’ reaction times, we found that participants choosing state 12 were indeed faster than participants choosing state 7 (*t*(122) = 3.05, *p* = .003, *d* = 0.57, *BF* = 11.9). Interestingly, participants choosing state 12 did not differ significantly in their reaction times from participants who chose state 11 that was predicted by model-free reinforcement learning (*t*(87) = 0.93, *p* = .35, *d* = 0.33, *BF* = 0.47), who in turn were also faster than participants choosing state 7 (*t*(51) = 2.51, *p* = .015, *d* = 0.91, *BF* = 3.6). Interestingly, participants choosing state 9 where as fast as participants choosing state 7 (*t*(82) = 0.43, *p* = .67, *d* = 0.09, *BF* = 0.03), which is somewhat surprising considering that the UVFA computation should be faster than the model-based computation, given that it is just a function approximator.

For our final exploratory analysis, we looked at the factors that predicted whether or not participants chose state 12 on the 101st trial. Because SF&GPI assumes that participants map previously learned policies onto novel tasks, this means that participants’ knowledge of the policy choosing state 12 should be predictive of the probability of choosing that state during the test trial. We therefore created a binary variable indicating whether or not participants chose state 12 on the 101st trial. We also created four independent variables. The first one was how often participants successfully chose state 12 during training and was expected to be highly predictive of choosing state 12 during the test trial, as predicted by SF&GPI. The second one was whether or not participants chose state 12 on the 100th trial, which we used to rule out a simple model-free repetition of the previous action. The third one was how often participants chose state 12 unsuccessfully (leading to negative rewards) during training, which we used to rule out simple experience effects. The final one was participants’ average reward during training, which we used to rule out simply using better strategies more generally. As predicted by SF&GPI, the frequency of successfully choosing state 12 during training was most predictive of choosing state 12 at test (standardized Bayesian Leave-One-Out cross-validation error LOO-*r*^2^ = 0.103). Choosing state 12 on the 100th trial (LOO-*r*^2^ = 0.086, *BF* = 269.2), the frequency of unsuccessfully choosing state 12 during training (LOO-*r*^2^ = 0.024, *BF* = 1.09 × 10^6^), as well as participants’ average rewards (LOO-*r*^2^ = 0.007, *BF* = 1.7 × 10^7^) were all less predictive of participants choosing state 12 on the 101st trial. These results provide further evidence that it was indeed participants’ knowledge of the policy choosing state 12 that helped them to do so on the 101st trial.

Summarizing the results of our preregistered Experiment 4, we found strong evidence for our pre-registered hypotheses that participants choose the state predicted by SF&GPI more frequently than any other state. Moreover, additional exploratory analyses showed that participants did not simply repeat previously successful choices at random, were faster due to the computationally less demanding SF&GPI strategy, and that their choice of the state predicted by SF&GPI was related to how well they had learned the reusable policy.

## Discussion

How do people learn to find rewards when they are confronted with multiple tasks? We studied participants’ behavior in a novel sequential decision task in which tasks varied as a function of feature-based rewards. We created tasks in which a participant’s decision during a final test trial discriminated between several models: two recent multi-task RL algorithms, as well as traditional model-based and model-free algorithms. Across four experiments, we found strong evidence for an algorithm that combines successor features with generalized policy iteration (SF&GPI). This suggests that people tackle new tasks by comparing policies of familiar tasks based on predictive state features.

There has been a recent surge of interest in multi-task RL in the machine learning community.^8–10^ The key question is what shared structure an agent can exploit across tasks to generalize efficiently. Different algorithms vary in their assumptions about this shared structure. We focused on two algorithms, UVFAs and SF&GPI, that exemplify particular classes of solutions. UVFAs exploit structure in the space of value functions, whereas SF&GPI exploits structure in task space.

Beyond the algorithms examined in our paper, there are a number of other important ideas that have been successful in machine learning. For example, hierarchical RL algorithms learn policy primitives that combine to produce solutions to different tasks,^11^ while meta-RL algorithms learn a learning algorithm that can adapt quickly to new tasks,^12, 13^ echoing the classic formation of task sets described by Harlow.^14^ Of particular note is the development of hybrids of UVFAs and SF&GPI known as universal successor features approximators, which combine the benefits of both approaches.^7^ Studying how people perform multi-task reinforcement learning can inform this line of research and help facilitate the development of algorithms with multi-task solving abilities on par with humans. We believe our study is an important initial step in that direction.

Multi-task RL has recently gained attention in computational neuroscience. Yang and colleagues^15^ trained a single recurrent neural network to solve multiple related tasks and observed the emergence of functionally specialized clusters of neurons, mixed selectivity neurons like those found in macaque prefrontal cortex, as well as compositional task representations reminiscent of hierarchical RL. Wang and colleagues^12^ proposed how meta-RL might be implemented in the brain, with dopamine signals gradually training a separate learning algorithm in the prefrontal cortex, which in turn can rapidly adapt to changing task demands.

There is also recent work showing that successor features can explain a wide range of puzzling phenomena related to the firing of hippocampal place cells and entorhinal grid cells,^16^ traditionally thought to encode the location of animals in physical and abstract spaces as a kind of cognitive map.^17^ Stachenfeld and colleagues proposed that, instead, these regions encode a predictive map, indicating which states the animal will visit in the future under a given policy (a special case of successor features in which each feature corresponds to a future state). Our work invites speculation that the predictive map may in fact be factorized into distinct state features, with different subsets of cells corresponding to different anticipated features of the environment. Furthermore, generalized policy improvement predicts that the activity of these cells should be tightly coupled to behavior on new tasks: how well the successor features for different policies are learned, and whether they are instantiated on the test trial, will govern which final states are considered, and ultimately which action is chosen. Finally, the successor features can be learned using a kind of vector-based temporal difference error,^5^ which quantifies the difference between the expected and the actually observed features. Such a vector-valued sensory prediction error may be encoded by dopamine neurons in the midbrain.^18^

One natural question that arises from our formulation of multi-task RL is where the feature weights **w** for the different tasks come from. One possibility is that they are imposed externally in explicit form, as they are in our study. They could also be learned from experience, potentially together with the low-dimensional feature representation *φ*.^19^ However, another possibility is that they are internally generated, more akin to feeling groggy or hungry as in our example from the introduction. The task defined by **w** would then correspond to the agent’s internal state, with each feature weight reflecting how valuable a given feature is based on the agent’s needs.^20^ These feature weights could be encoded as part of a generic latent state representation, such as the one thought to be encoded in orbitofrontal cortex,^21, 22^ or in brain regions specific to representing physiological needs, such as the hypothalamus^23^ or the insula.^24^ Such a perspective can help resolve the “reward paradox”,^25^ a key challenge of applying RL as a theory of human and animal learning, which typically assumes an external reward function that does not exist in natural environments. This view predicts that inducing different motivational states (for example, hunger, thirst, sleepiness) would correspond to naturalistically varying the feature weights **w**.

Our study has several limitations. Most notably, our definition of multi-task RL is fairly narrow: we consider tasks that share the same deterministic state transition function and only differ in the reward function. Furthermore, our design is restricted to the tabular case, with a discrete enumerable state space. Constraining our study in this way allowed for a clean test of our hypotheses, yet as a result it falls short of capturing the full plethora of multi-task RL behaviors, such as compositionality of tasks and policies,^26, 27^ or discovery of shared task representations.^28^ Indeed, the overly simplistic design could explain why we found no evidence for UVFAs: even though there is smoothness in the task space, there is no smoothness in the state space, which might be crucial for humans to leverage this type of generalization. Future studies could investigate human multi-task RL in richer, more complex domains, such as video games.^29^

Finally, our experiment was designed to distinguish alternative accounts of multi-task RL only based on performance on a test trial. From this vantage point, our treatment of the multi-task RL problem might seem unsatisfactory, since it only speaks to how values and policies are transferred, rather than how they are learned in the first place. This limitation stems from our goal to distinguish between entire classes of algorithms, without committing strongly to particular instantiations from each class. There are many different ways to learn the values of UVFAs or the policies and successor features of SF&GPI, resulting in different predictions about the process of learning that are secondary to the multi-task problem itself. By focusing on predictions that are invariant to the particular learning algorithm, and preregistering our predictions for the final experiment, we were able to maximally distinguish between algorithms, thus narrowing down the space of viable models, and paving the way for future studies of learning.

The ability to flexibly adapt to changing task demands and creatively use past solutions in new situations is a hallmark of human intelligence that is yet to be matched by artificial agents.^2^ In the current study, we investigated how humans accomplish this behavior in the framework of multi-task RL. Using a sequential decision making task, we found evidence that people transfer knowledge from familiar tasks to unfamiliar tasks by comparing previous solutions and choosing the one which leads to states that are most favorable according to the new task. This strategy offers an efficient alternative to standard model-free and model-based approaches. We believe that studying how humans learn and search for rewards across multiple tasks will allow our models to generalize to increasingly broad and complex domains.

## Methods

### Computational Models

In this section, we describe the general theoretical setup that motivates the RL models whose predictions we tested experimentally. A complete description of the models can be found in the Supporting Information.

We define the multi-task RL problem using the formalism of Markov decision processes (MDPs). An MDP is defined as a tuple 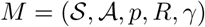, where 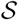 and 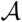 are the state and action spaces, *p*(*s′*|*s, a*) is the probability of transition to state *s′* after taking action *a* in state *s*, *R*(*s*) is a random variable that represents the reward received upon transitioning into state *s*, and *γ* ∈ [0, 1) is a discount factor that down-weights future rewards as an exponential function of their temporal distance in the future.

To apply this formalism to the multi-task learning scenario, it is useful to introduce a set of *features φ*(*s′*). Returning to the example from the introduction, if the feature dimensions are food, coffee, and atmosphere, then the features might be *φ*(burger joint) = [90, 10, 50], *φ*(coffee shop) = [10, 90, 90], and *φ*(diner) = [50, 50, 50]. That is, the burger joint has good food, bad coffee, and a decent atmosphere; the coffee shop has bad food, good coffee, and a good atmosphere; and the diner has decent food, decent coffee, and a decent atmosphere.

We assume the agent faces a sequence of tasks, with each task defined by a vector of weights **w**. The weights **w** determine how valuable each feature is, thus inducing a task-specific reward function *R*_**w**_. The expected one-step reward when transitioning from *s* to *s′* can be computed as the sum of the encountered features *φ*, weighted by the task-specific feature weights **w**:

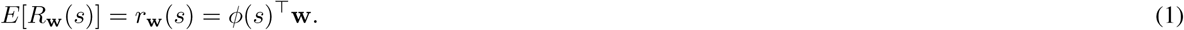

In our running example, the task to solve when feeling hungry can be expressed as **w**_hungry_ = [1, 0, 0]. In that case, going to the burger joint would yield an expected reward of *φ*(burger joint)^⊤^**w**_hungry_ = [90, 10, 50]^⊤^[1, 0, 0] = 90, while going to the coffee shop would yield a reward of merely *φ*(coffee shop)^⊤^**w**_hungry_ = [10, 90, 90]^⊤^[1, 0, 0] = 10. Similarly, the task to solve when feeling groggy can be expressed as **w**_groggy_ = [0, 1, 0], **w**_hungry_and_groggy_ = [1, 1, 0], and so forth.

This means that instead of solving a single MDP, the agent must solve a *set* of MDPs that share the same structure but differ in their reward functions (note that this only one specific class of multi-task RL problems; more generally the MDPs can differ in more than just the reward function). Specifically, the problem facing the agent is to find a policy *π*_**w**_: *S → A* that maximizes the discounted future rewards 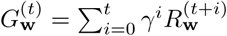, where 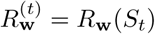 is the reward received at time *t*. The action-value function of a policy *π* on a particular task **w** can be defined as:

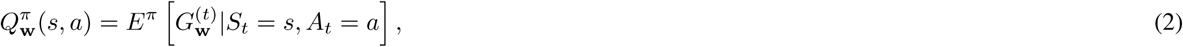

where *E*^*π*^[⋅] denotes the expected value when following policy *π*.

One of the benefits of learning about multiple tasks at the same time is the possibility of transferring knowledge across tasks with little new learning.^8, 30^ There are two sources of structure an agent can exploit in the service of transfer: the similarity between the solutions of the tasks (either in the policy or in the associated value-function space), or the shared dynamics of the environment. Here, we compare two classes of multi-task RL models, which directly tap into these two sources of structure.

*Universal value function approximators* (UVFAs) extend the notion of value functions to also include the description of a task, thus directly exploiting the common structure in the associated optimal value functions.^4^ The basic insight behind UVFAs is to note that the concept of an optimal value function can be extended to include a description of the task as an argument. One way to do so is to specify a “universal” optimal value *Q*^∗^(*s, a*, **w**) as a function of task **w**. The agent learns an approximation 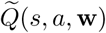. A sufficiently expressive UVFA can identify and exploit structure in the value functions across tasks, and thereby efficiently generalize to novel tasks.

*Successor features and generalized policy improvement* (SF&GPI) exploits the common structure in the environment and capitalises on the power of dynamic programming.^5, 6^ The SF&GPI approach is based on two concepts: successor features and generalized policy improvement. The successor features of a state-action pair (*s, a*) under policy *π* are given by

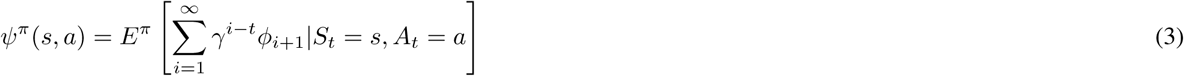

Intuitively, successor features correspond to the cumulative features that the agent can expect to see over the long run when following a given policy *π*. Successor features allow for the immediate calculation of a policy’s values on any task **w**.^31, 32^ Moreover, since they satisfy the Bellman equation, successor features can be learned by standard reinforcement learning algorithms. Generalized policy improvement (a generalization of the classic policy improvement algorithm) computes a policy based on a set of value functions rather than on a single one. If an agent has learned the successor features 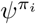 for policies *π*_1_, *π*_2_, …, *π*_*n*_ and is confronted with a new task **w**, then it can easily compute the values of the policies on the new task as 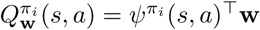, which can be used to find the best suitable policy for the current task, 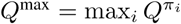.

Intuitively, SF&GPI corresponds to learning the features of states as well as policies that have resulted in high rewards in previous tasks. If a novel task appears, then given the learned features and transition structure, the agent can evaluate what the best past policy is to apply to the current task. SF&GPI is appealing because it allows transfer to take place between any two tasks, regardless of their temporal order. SF&GPI is also closely related to recent applications of successor features to modeling behavioral^33, 34^ and brain data,^35^ as well as to findings showing that participants reuse previously discovered solutions to solve new but related tasks.^36, 37^

UVFAs and SF&GPI generalize to new tasks in two different ways. Whereas the UVFAs aim to generalize across the space of tasks by exploiting structure in the underlying value function, SF&GPI aims to exploit the structure of the reinforcement learning problem itself.^7^

## Experimental Methods

### Participants

Participants were recruited from Amazon Mechanical Turk for Experiment 1 (*N* = 226, 92 female; mean±s.d. age: 35.2±10.3 years), for Experiment 2 (*N* = 202, 85 female; mean±s.d. age: 37.6±13.3 years), for Experiment 3 (*N* = 200, 89 female; mean±s.d. age: 35.8±11.2 years) and for Experiment 4 (*N* = 500, 214 females; mean±s.d. age: 34.66 ± 9.11). In all of the experiments, participants were paid a participation fee of US$1.50 and a performance contingent bonus of up to US$2.00. Participants earned on average US$2.14±0.13 and spent 32.2±3min on the task in Experiment 1, earned US$2.64±0.20 and spent 29.9±4min in Experiment 2, earned US$2.53±0.15 and spent 34.3±5min in Experiment 3 and earned US$2.58±0.28 and spent 30.8±5min in Experiment 4. Participants were only allowed to participate in one of the experiments and were required to have a 90% human interaction task (HIT) approval rate and 100 previously completed HITs. No statistical methods were used to pre-determine the sample sizes for Experiments 1-3, but our sample sizes are similar to or larger than those reported in previous publications.^33, 38, 39^ For the preregistered Experiment 4, we aimed to collect 500 participants in total, making this sample size more than twice as large as in our previous studies. The Harvard Internal Review Board approved the methodology and all participants consented to participation through an online consent form at the beginning of the survey.

### Design

All participants went through the same training task with training weights **w**_train_ = {[1, −1, 0], [−1, 1, 0], [1, −2, 0], [−2, 1, 0]} and a test weight **w**_test_ = [1, 1, 1]. Participants encountered each training weight on 25 randomly chosen trials of 100 trials in total. On the 101st trial, the novel test weight was introduced. The task structure was a two-step decision tree with three nodes on each level and one node at the start, leading to 13 states in total. The transitions between states were deterministic, which means that the same choice in the same state always led to the exact same next state. The final states contained three feature values, which we adapted to create diverging model predictions. The feature values for all states are shown in Tab. 1 for all experiments.

**Table 1.** Feature values used for all Experiments. The value shown for each state represent the values *φ*(state) which participants encountered when ending up in this state.

| State | Experiment 1 | Experiment 2 | Experiment 3-4 |
| --- | --- | --- | --- |
| 5 | [0, 10, 0] | [10, 20, 90] | [0, 10, 0] |
| 6 | [100, 0, 0] | [100, 0, 20] | [100, 0, 0] |
| 7 | [70, 70, 70] | [70, 70, 170] | [40, 40, 130] |
| 8 | [90, 0, 0] | [90, 0, 40] | [90, 0, 0] |
| 9 | [100, 100, 0] | [150, 150, 0] | [100, 100, 0] |
| 10 | [0, 90, 0] | [0, 90, 30] | [0, 90, 0] |
| 11 | [0, 0, 10] | [30, 10, 50] | [0, 0, 10] |
| 12 | [0, 100, 60] | [0, 100, 160] | [0, 100, 60] |
| 13 | [10, 0, 0] | [20, 10, 100] | [10, 0, 0] |

The reward on each trial was calculated by multiplying the features of the final state with the current task weights **w**. The detailed pre-registration for Experiment 4 can be accessed at https://osf.io/cuxqn/.

### Materials and procedure

Participants played a game in which they were a tradesperson scavenging a medieval castle for resources to trade. Additionally, they were told that there were three types of resources (wood, stone, and iron) that they could sell for different prices. Moreover, some resources could not be sold but instead needed to be disposed of at a cost. The prices and costs for the different resources were shown to participants on every trial, with prices presented as positive and costs as negative money. Furthermore, participants were told that they would see the daily market price on every trial before they entered the castle. Participants initialized a trial by entering the castle through pressing the space key. Once the trial started, they saw three different doors, which they could select by pressing 1, 2 or 3 on their keyboard. After they had chosen one of the three doors and walked through it, they were in a new room which always again had three different doors that they could choose by pressing 1, 2 or 3. After their second decision, they entered a final room in which they found resources (corresponding to *φ*(*s′*)) which were then multiplied by the costs/prices of that day and converted to USD by dividing the trial score by 100. Every specific path always corresponded to the same chosen doors, where we assigned different pictures of doors to the choices at random for each participant before the experiment started. Once the experiment started, everything was deterministic, with the same choices always leading to the same rooms and all of the final nodes always having the exact same feature values. The only thing that changed over trials was the task defined by the weight **w**. On each trial, the current prices/costs for every resources as well as their decision history (i.e. which door they had chosen on the previous step) was shown to participants on top of the screen. Participants’ bonus was determined by randomly sampling one of the 101 trials and using that trial’s payoff.

### Statistical Tests

All reported binomial tests are calculated based on exact testing against the null hypotheses of chance *p* = 1/9. We additionally report Bayes factors (*BF*) for all tests. A Bayes Factor quantifies the likelihood of the data under *H*_*A*_ relative to the likelihood of the data under *H*_0_. For all tests based on proportions, we tested the null hypothesis that the probability of a choosing a final state is *p*_0_ against alternative that it is *λ* ~ logistic(*λ*_0_, *r*), with *λ*_0_ = logit(*p*_0_) and *λ* = logit(*p*). The parameter *r* is a scaling factor and set to 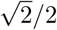. We computed the Bayes factor via Gaussian quadrature and drew posterior samples via independence Metropolis-Hastings.^40^ For comparing participants’ reaction times in Experiment 4, we used the Bayesian t-test for independent samples,^41^ with the Jeffreys-Zellner-Siow prior and scale set to 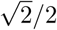. Finally, we used Bayesian multi-level logistic regressions for Experiment 4 with a prior on the coefficients of *β* ~ *N* (0, 10)) and approximated the Bayes Factor using bridge sampling.^42^

## Code Availability

Code for all models and analyses is available at https://github.com/tomov/MTRL.

## Data Availability

Anonymized participant data and model simulation data are available at https://github.com/tomov/MTRL.

## Acknowledgements

We thank Nick Franklin and Wanqian Yang for helpful discussions. This research was supported by the Toyota corporation, the Office of Naval Research (Award N000141712984), and the Harvard Data Science Initiative.

## Author contributions statement

M.T. and E.S. contributed equally. M.T., E.S. and S.G. conceived the experiments, M.T. conducted the experiments, M.T. and E.S. analysed the results. All authors wrote the manuscript.

## Supporting Information

### Model Specifications

We formally specify all models below. Even though it is possible to change specific parts of the models’ implementations, all of our predicted effects are largely independent of implementational details.

For all models, we evaluated performance on the test trial using a softmax policy based on the learned Q-values, with inverse temperature *β* = 10. Since we were primarily interested in comparing models based on asymptotic performance, this ensured that the models were placed on an equal footing, even though they were trained using different methods.

#### Model-free learning

We trained a model-free agent on the same training tasks and environment as the participants. We used tabular Q-learning:^3^ after taking action *a* and transitioning from state *s* to state *s′*, the Q-value for state-action pair (*s, a*) was updated as:

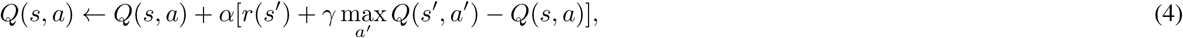

where *r*(*s*) = *φ*(*s*)^⊤^**w** for training task **w**. The learning rate was *α* = 0.1 and the discount factor was *γ* = 0.99. Values were mapped to choices using an *E*-greedy exploration strategy with *ϵ* = 0.9:

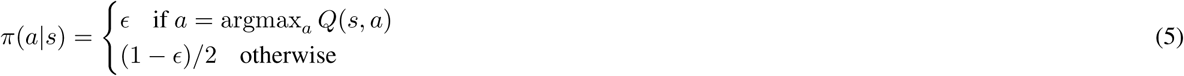

Q-values were initialized randomly between 0 and 0.00001 to break ties initially. Each training task was encountered for 200 training episodes (trials). Note that this approach ignores the features *φ* and task weights **w** and only considers states, actions, and the final reward.

#### Model-based learning

We assume the model-based agent has perfect knowledge of the environment. We computed the Q-values for each task **w** separately using value iteration:^3^ the value of each state *s* was updated according to:

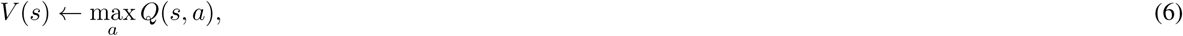

where

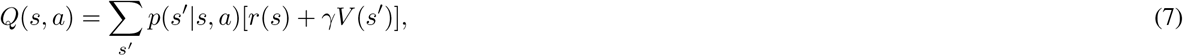

where *r*(*s*) = *φ*(*s*)^⊤^**w** for task **w**. This update was repeated for all states until convergence, defined as the largest update being less than 0.01. Note that, unlike model-free learning, value iteration was also applied to the test task.

#### Universal value function approximators

One way to train a UVFA online involves using a deep Q-learning network. Since we were interested in asymptotic performance, we instead trained it in a supervised way.^4^ For each training task, we computed the Q-values using value iteration as described in the model-based section. We then trained a 3-layer feedforward function fitting neural network (fitnet in MATLAB) with 10 hidden units to predict the Q-values for each training task. The transitions between states were deterministic, *Q*(*s, a*) = *V* (*s′*), where *p*(*s′*|*s, a*) = 1, so for inputs to the network we used (**s**, **w**)-tuples, where **s** is a one-hot encoding of the state and **w** is the task weights vector. Each training tuple and the corresponding Q-value were repeated 100 times. The Q-values for the test task were generated as the predictions of the network for (**s**, **w**_test_).

#### Successor features and generalized policy improvement

We assume the agent learns a policy and its corresponding successor features for each training task.^7^ While these could be computed online using temporal difference learning, we were primarily interested in asymptotic performance, so we computed the optimal policy for each training task using value iteration, as described in model-based section. We then computed the successor features for each policy *π* using dynamic programming by iteratively applying the Bellman equation until convergence (maximum change of less than 0.01):

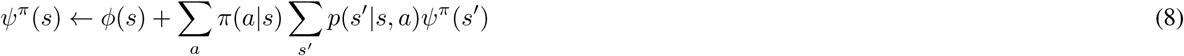

We computed the Q-values on the test task **w** using generalized policy improvement. This involved iterating over the policies and computing the expected reward of the test task **w**, using the successor features for each policy *π*:

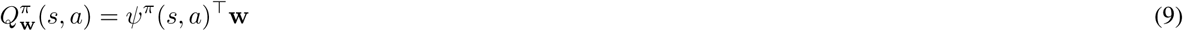

The Q-value for each state-action pair was then chosen based on the policy that will perform best on the test task **w**:

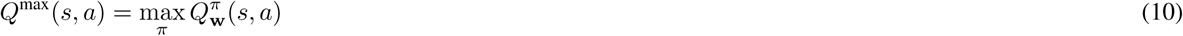

### Effect of exclusion criterion

We assess the effect of exclusion criteria onto our main finding of participants choosing the SF&GPI state more frequently than what was expected under the chance level of *p* = 1/9. First, we assess the proportions of people choosing the SF&GPI state without any exclusion of participants. Doing so, we find that 40 out 226 participants choose state 12 in Experiment 1 (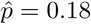, exact binomial test: *p* = .003, *BF* = 12.2), 48 out of 202 participants chose state 12 in Experiment 2 (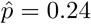, exact binomial test: *p* = 4.22 × 10^*−*7^, *BF* = 38408), and 42 out of 200 participants chose state 12 in Experiment 3 (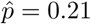, exact binomial test: *p* = 6.21 × 10^*−*6^, *BF* = 434). Next, we assess what happens to our results if we exclude participants who achieved an average reward below the median during the training trials. Doing so, we find that 21 out of 113 participants chose state 12 in Experiment 1 (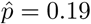, exact binomial test: *p* = .016, *BF* = 3.3), 30 out of 106 participants chose state 12 in Experiment 2 (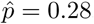, exact binomial test: *p* = 1.17 × 10^*−*5^, *BF* = 1376) and 31 out of 96 participants chose state 12 in Experiment 3 (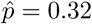, exact binomial test: *p* = 2.42 × 10^*−*8^, *BF* = 3.39 × 10^5^). We therefore conclude that our main effect is largely independent of the chosen method of participant exclusion.

